# Red Wood Ants prefer mature Pine Forests in Variscan Granite Environments (*Formica rufa*-group)

**DOI:** 10.1101/2020.11.26.390112

**Authors:** Gabriele M. Berberich, Martin B. Berberich, Matthias Gibhardt

## Abstract

We used presence/absence data of 5,160 red wood ant nests (RWA; *Formica polyctena*) acquired in a systematic large-scale area-wide survey in two study areas (≈350 ha) in the Oberpfalz, NE Bavaria, Germany to explore for the first time the influence of variable (e.g., forest type, tree age) and quasi-invariant factors (e.g., tectonics, geochemical composition of the bedrock) on nest size, spatial distribution and nest density for Variscan granites. A combination of the forest type (mature pine-dominated forests (≥80–140 years) as main variable factor and the geochemical property of the Variscan granites with their high natural Radon potential and moderate heat production as main quasi-invariant factor could explain the high nest numbers in both study areas. In addition, the spatially clustered distribution patterns of the observed nests suggest a strong interaction between nests and their quasi-invariant environment, especially the directionality of the present-day stress field and the direction of the tectonically formed “Erbendorfer Line”. In general, such a combination of variable and quasi-invariant factors can be addressed as particularly favorable RWA habitats.

## 1 Introduction

Geological and tectonic processes are not only fundamental driving forces, e.g., for the regional architecture of rocks, and evolution of fault systems, but also of long-term global biodiversity patterns and a wide range of environments and ecosystems due to tectonic shifts in the arrangement of the continental crust (Eisbacher 1991; Valentine & Moores 1970; Descombes et al. 2017). Understanding the processes that govern biodiversity, the relationship of organisms to specific habitats, their interspecific relationships as well as their distribution and occurrence are central questions of biology and ecology.

Ants, a geographically widespread taxon (e.g., Hölldobler & Wilson 1990) have major ecological impacts in terrestrial ecosystems. Especially, red wood ants (RWA; *Formica rufa-*group s. str.; Hölldobler & Wilson 2010), which form very large, often polydomous colonies (e.g., Ellis & Robinson 2014) are important ecological keystone species (e.g., Frouz & Jílková 2008). As important ecosystem engineers, RWA contribute to the diversity and formation of a dynamic balance in forest habitats (Wellenstein 1990), influence the circulation and the distribution of nutrients (Klimetzek & Kaiser 1995), regulate insect pests as biological control agents (e.g., Robinson et al. 2016) and contribute to soil-forming processes (Frouz et al. 2008). In entomological studies, the spatial occurrence and distribution of RWA had been mostly attributed to specific forest factors, e.g., canopy cover and edge (e.g., Risch et al. 2008), fragmentation (Punttila & Kilpelainen 2009), tree species, characteristics and age (e.g., Gibb et al. 2016), food supply (e.g., Iakovlev et al. 2017), social organization of species (Ellis & Robinson 2014), but also abiotic factors such as solar radiation (Kadochová & Frouz 2014), and altitude (Vandegehuchte et al. 2017).

Recent studies, which investigated a combination of geoscientific and biological factors (also known as “GeoBio-Interactions”), developed RWA as biological indicators for otherwise undetected tectonic activity and showed that RWA nests were eight times more likely to be found within 60 m of known tectonic faults (Berberich et al. 2016a; Del Toro et al. 2017; Berberich et al. 2019). Furthermore, geogenic gases (e.g. CO_2_, Helium, Radon; Berberich et al. 2019), fault-related emissions of CH_4_ (–37 % ^13^C-CH_4_ signature; Berberich et al. 2018a), volatile organo-halogens (CH_3_Cl, CHCl_3_, CHBr_3_), as well as alkanes and limonene (Berberich et al. 2016a) play a decisive role in the settlement of RWA nests.

A combined analysis of specifically variable (biotic) factors and quasi-invariant (geo-tectonic) factors in two comparable study areas Falkenberg (FB) and Münchsgrün (MG), both located in the Oberpfålzer Wald region (NE Bavaria, Germany), was considered appropriate to further investigate GeoBio-Interactions in Variscan Granite environments. To date, no systematic large-scale survey of RWA nests is available for both study areas. In addition, literature and geological maps contain only incomplete or no information on tectonic fault systems for both study areas.

In this study, we addressed for the first time, forest-tectonic interactions based on presence/absence data of RWA nests for Variscan Granites by asking three interrelated main questions: (1) What influence do variable factors such as forest type and tree age have, and (2) What influence do quasi-invariant factors, such as tectonics, geochemical composition of the underlying bedrock have on a) nest size, b) spatial distribution of RWA nests, and c) nest density, and (3) Is there an explanation of the extremely high RWA nest number and density in both study areas? We have asked these questions specifically with regard to individual RWA nests. It is expected that the results will further improve and complement the understanding of the “GeoBio-Interactions”, which will be applicable for further investigations of RWA. Additionally, the database of RWA nest sizes and locations will be a tool for the future forest management with regard to RWA protection in both study areas.

## 2 Materials and methods

### 2.1 Location, forest types and tectonic settings

#### 2.1.1 Location

The two densely forested study areas, Münchsgrün (MG) and Falkenberg (FB), characterized by a continuous low mountain range, were located in the Oberpfälzer Lake district (Tirschenreuth county), approx. 50 km East of the city of Bayreuth and 20 km West of the Czech Border (NE Bavaria, Germany; Fig. 1). The valleys were used for agriculture, settlements and transport routes (Glaser et al. 2007). MG study area (475–530 m), a smoothly NE-SW inclined terrain, was located between the city Mitterteich (5 km North) and the city Tirschenreuth (5 km Southeast). FB study area (470– 545 m), a partially rugged terrain, was located between the cities of Falkenberg (2 km East) and Tirschenreuth (4 km West). The region, also called “Siberia of Bavaria”, is characterized by a low annual average temperature of < 7 °C, a short vegetation period (< 140 days), 30 icy days/year and a high precipitation rate (average 700–800 mm/year; LK-Tir. 2020).

**Fig 1.**
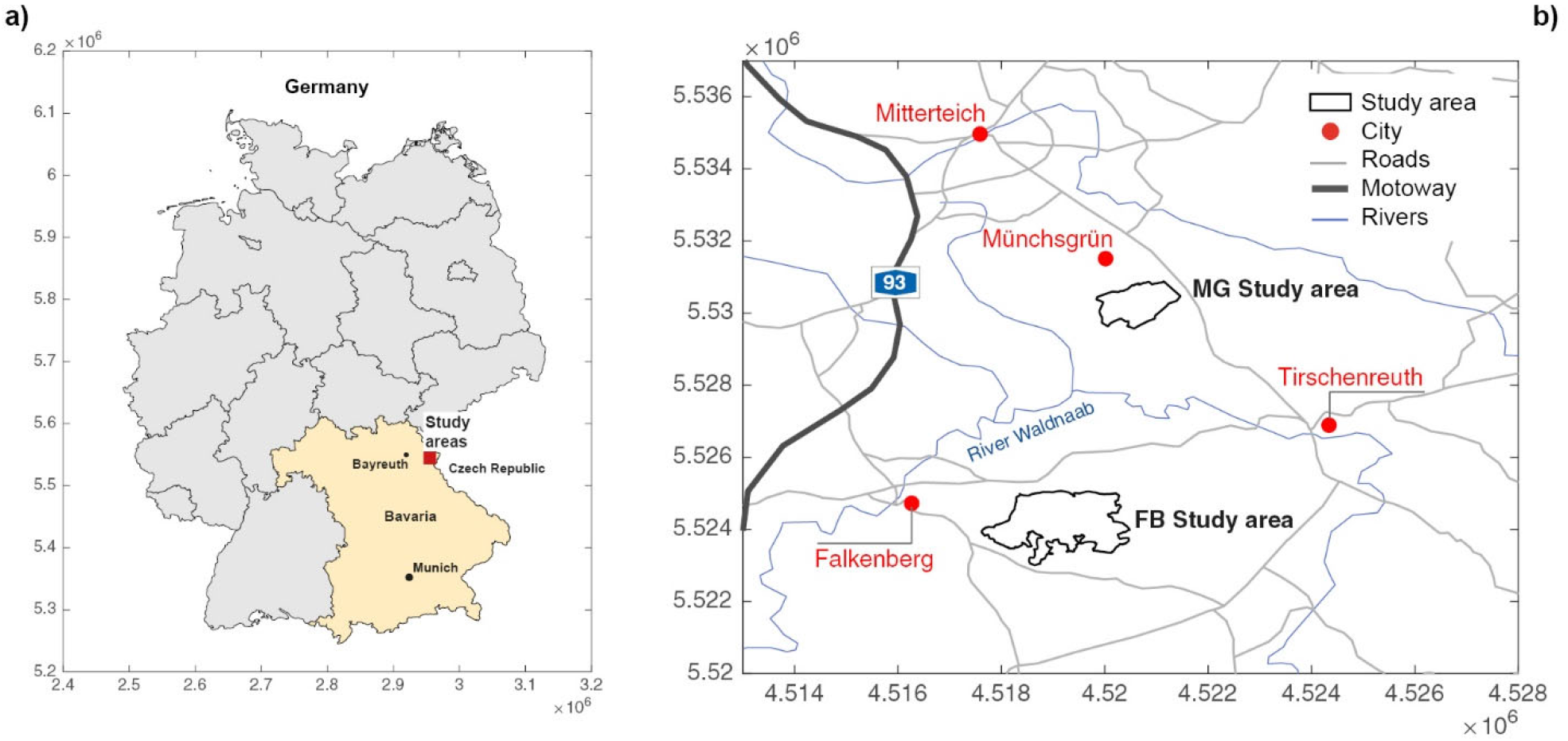
a) Position of both study areas within Germany close to the Czech border; b) detailing location in the Oberpfälzer Lake district in Tirschenreuth county, NE Bavaria.

#### 2.1.2 Forest types

Both study areas were located in managed forests, with an integrative sustainable forest management, which was administered by the Bavarian State Forest (BSF), Regensburg. The objective of this management approach is to establish a “permanent forest”, i.e. a vertically structured, ecologically valuable forest with different trees ages and species, which is stocked with the best possible quality of stand, and in which the natural self-regulating mechanisms are used and maintained. Furthermore, the protective and recreational functions of the forest and its function as a source of raw materials are promoted. The forest managed by the BSF in MG (MG_BSF_) consisted of only one, in FB (FB_BSF_) of eight forest sections: 1= Tiefe Lohe; not mapped in this study; 2= Alter Forstmeister; 3= Himmelreich; 4= Ebene; 5= Steinlohe; 6= Weißgärberwiese; 7= Spechtnerbrückl; 8= Winterleite (Fig. 2; BSF 2009).

**Fig 2.**
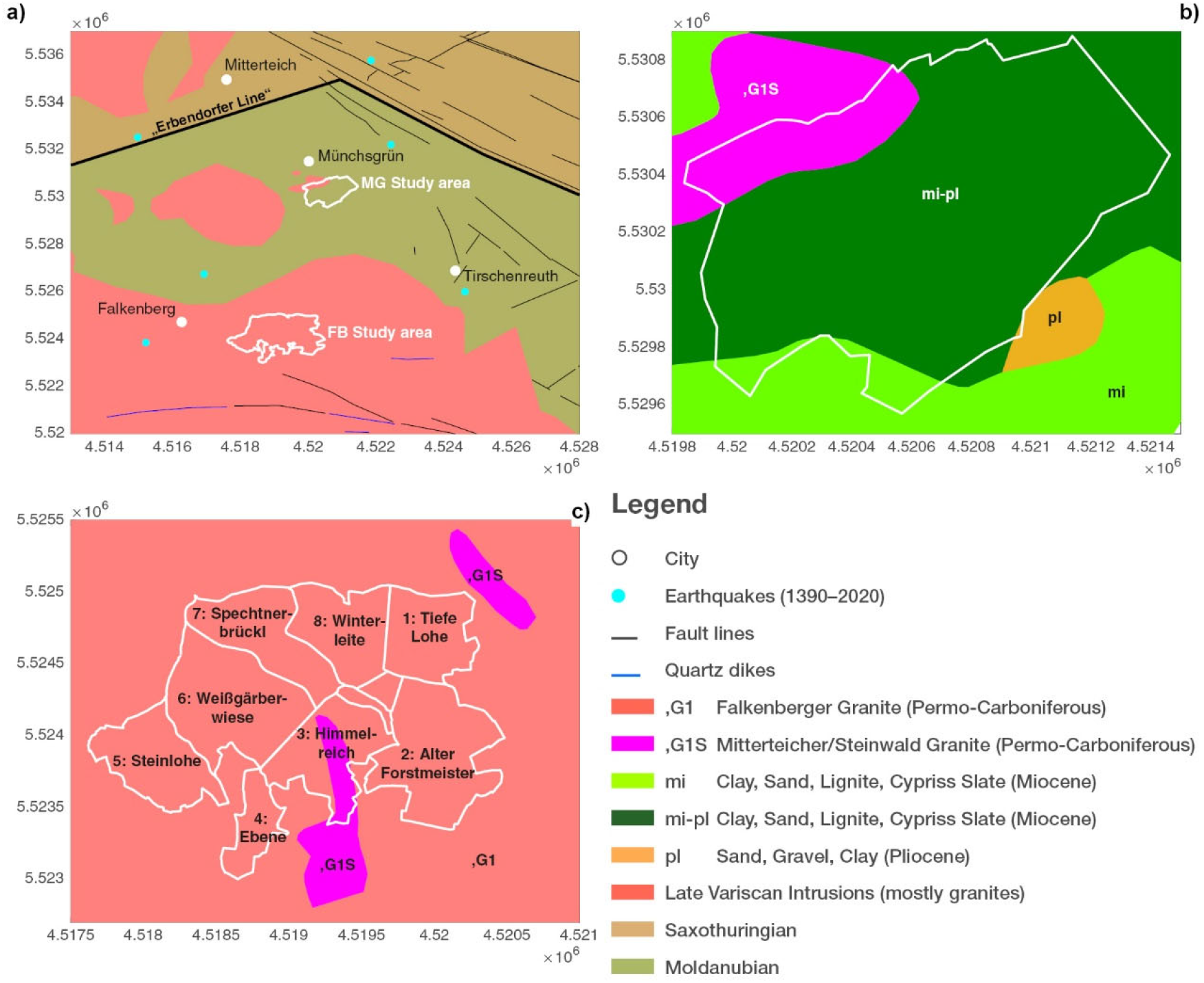
Tectonic setting of both study areas with (a) major tectonic units, faults (black lines), earthquake events (blue dots) taken from literature (see list under reference section geological maps), b) detailed geologic setting of the Münchsgrün (MG), and c) Falkenberg (FB) study areas with FB forest sections (1-8).

In both study areas, the coniferous forest was strongly-dominated by two primary tree species (TS_prime_): spruce (*Picea abies*; FB_BSF_ ≈50 %; MG_BSF_ ≈40 %) or pine (*Pinus sylvestris*; FB_BSF_ ≈50 %; MG_BSF_ ≈56 %; BSF 2009). Deciduous trees such as black alder (*Alnus glutinosa*), hornbeam (*Carpinus betulus*), red oak (*Quercus ruba*) or European white birch (*Betula pendula*) accounted for less than ≈2 % as TS_prime_. Mixed coniferous stands were characterized by pine (TS_prime_) and spruce (TS_sec;_ secondary tree species) and amounted to ≈80 % (FB_BSF_) and ≈95 % (MG_BSF_). Furthermore, mixed coniferous-deciduous stands in FB_BSF_ consisted of pine and spruce (TS_prime_) and beech (TS_sec_; up to 10 %); in the MG_BSF_ of pine and birch (TS_prime_) and spruce (TS_sec_; ≈58 %) (BSF 2009; Tab. 1).

**Table 1.**
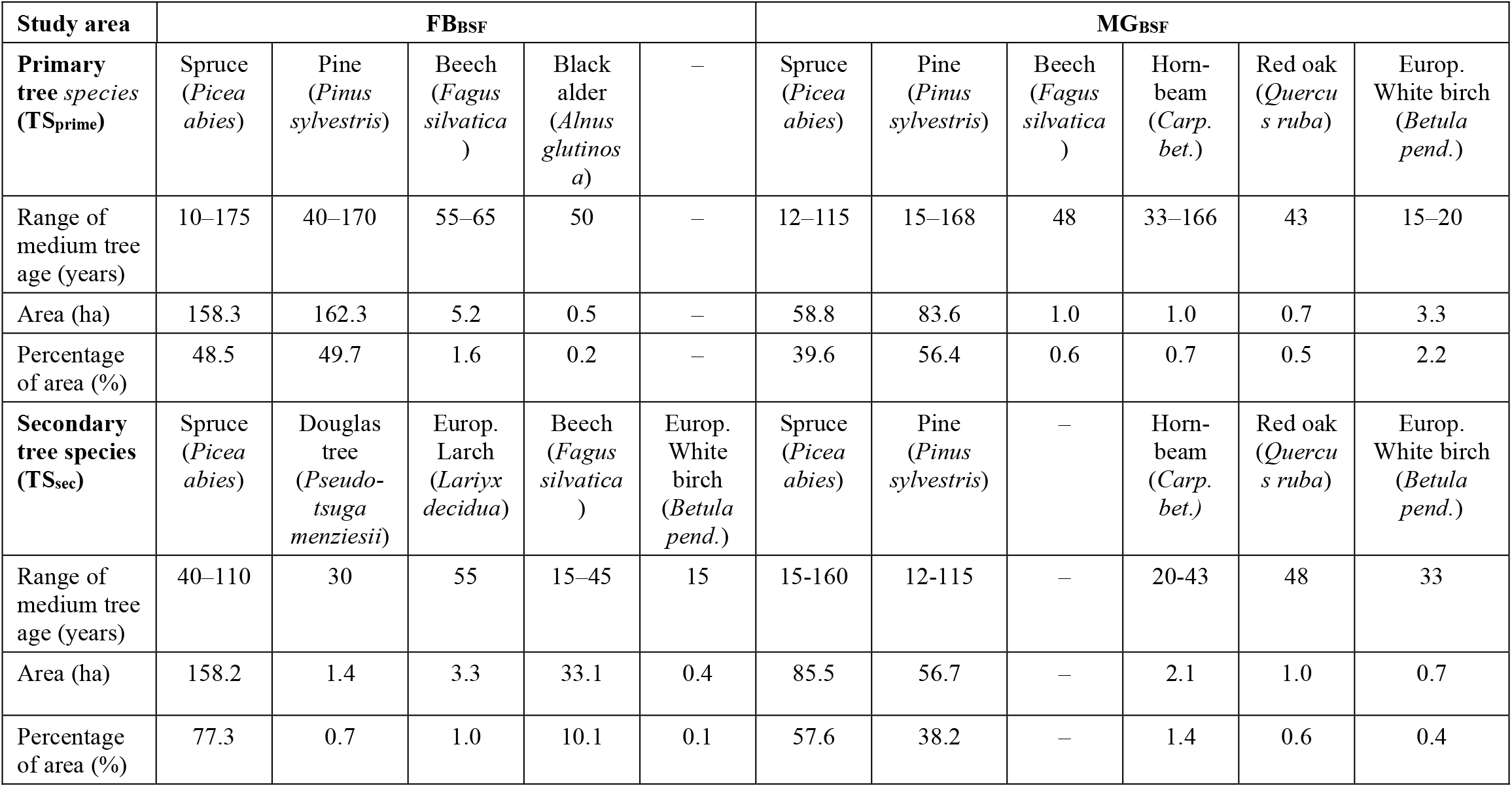
Primary tree species, medium age and area size of primary tree species (TS_prime_) and secondary tree species (TS_sec_) in MG_BSF_ and FB_BSF_ study area (BSF 2009); – = not present.

#### 2.1.3 Geologic and tectonic setting

Complex tectonic, magmatic and geologic processes that have occurred since the Paleozoic, characterize the region of the two study areas in the North-East Oberpfalz (“Upper Palatinate”). The most important tectonic processes were collision of microplates with the continent Gondwana, rifting, volcanism, thrusting, subduction, and uplift. The formation of the Variscan Orogen began at the end of the Devonian Period (≈360 Mio. years ago) as a result of a continental-continental collision. At the end of the Carboniferous (≈300 Mio. years ago), the Variscan Orogen began to collapse, accompanied by mantle-induced crustal extension and magmatic activity, which led to granite intrusions (Scharfenberg & De Wall 2016). During the Alpine orogeny (100–30 Mio. years ago), compression and transpression tectonics, rifting (e.g., Eger rift system), volcanism, subsidence and uplift, and the formation of Neogene sedimentary basins (e.g., Mitterteicher Basin) prevailed (e.g., Glaser et al. 2007; Peterek & Schunk 2009).

Today, the Oberpfalz region is affected by a present-day compressional stress field oriented in NW-SE to NNW-SSE direction (Heidbach et al. 2016). A compilation of faults from several publications showed that NW-SE trending fault systems predominate; W-E, WNW-ESE and NE-SW trending faults are also present (Fig. 2a). Quartz dikes are arranged in dominant NW-SE directions; some of them show W-E, WNW-ESE directions (e.g., Emmert et al. 1981; Peterek & Schunk 2008; LfU 2015). Two different basement units, which delimit regions with different tectono-metamorphic history (Rohrmüller et al. 2000) characterize the regional tectonics: the Saxo-Thuringian unit in the North, and the Moldanubian unit in the South, which underlie both study areas. Both units are delimited by a large, fault system several tenth of kilometers long, the “Erbendorfer Line” which runs in NE-SW and NW-SE direction (e.g., Hofmann 2003; Glaser et al. 2007; Galadí et al. 2009).

The oldest geological formation in both study areas is the Permo-Carboniferous crystalline basement, that is composed of granitic intrusions that form the Younger Intrusive Complex (YIC; Scharfenberg & De Wall 2016). In the MG study area, the “Mitterteicher Granite” (also known as Steinwald granite, depending on the location, henceforth Mitterteicher/Steinwald granite; 312–310 Mio. years), is a medium-grained ± porphyric Muscovite-Monzogranite. This granite is covered up to ≈89 % by Oligocene-Pliocene clastic sediments (clay, silt, sand and gravel) as part of the “Mitterteicher Basin” fillings (Peterek & Schunk 2009) and small lenses of Pleistocene-Holocene valley fillings (loam and sand; Fig. 2b). Two different granites of the YIC are present in the FB study area: a) the younger, 1.2 km long, lobate-shaped (10–400 m wide), N-S running Mitterteicher/Steinwald granite intrusion in the South which separated the forest section 3: Himmelreich into three parts, and b) the older “Falkenberger Granite” (coarse-grained porphyric Andalusite-Sillimanite-Monzogranite; ≈315 Mio. years) which made-up the rest of the study area (Fig. 2c; LfU 2013; Scharfenberg & De Wall 2016).

In both study areas, soil, vegetation (forest stands, agriculture), and sediment cover left only sparse outcrops, which led to a limited and/or incomplete knowledge of the entire tectonic regime. Sporadic weak to moderate earthquakes, which mostly occur in a shallow crustal depth (< 20 km) and whose local magnitudes (M_L_, Richter scale) rarely exceed 3.5, are mostly concentrated in the NW of the city of Mitterteich (LMU 2020).

### 2.2 Mapping and data collection

Approximately, ≈130 ha (MG_BSF_) and ≈170 ha (FB_BSF_) had been mapped in the Bavarian State Forest (BSF). Due to the high abundance of RWA nests in this forest, some adjacent areas were also mapped, so that the total mapped area amounts to ≈150 ha (MG_tot_) and ≈200 ha (FB_tot;_ Fig. 3). The mapping of RWA nests followed the approach already described in previous publications (Berberich et al. 2016a). In both study areas, approximately 5,160 inhabited RWA nests were mapped area-wide (MG: October 2015 and April 2016; FB: May and October 2019), with GPS receivers (Garmin 60CSx/62S). In the field, six nest height classes (NH; start-ups: 0.01–0.10 m, short: 0.11–0.50 m, medium: 0.51–1.00 m, tall: 1.01–1.50 m, very tall: 1.51–2.00 m, extra tall: >2.00 m) and five diameter classes (ND; small: 0.01–0.50 m, medium: 0.51–1.00 m, large: 1.01–1.50 m, very large: 1.50–2.00 m, and extra-large > 2.00 m) and the nest location (e.g. within the forest, forest roads, forest edges), were classified (*c*.*f*. Berberich et al. 2016a, 2016c). In addition, information on the herb layer on/around the RWA nest, e.g., nettles, black berry, gras, were collected. Furthermore, the visible nest material was identified and classified into three classes in the field: only spruce needles (100 %), only pine needles (100 %) or ≈50 % spruce and ≈50 % pine needles.

**Fig 3.**
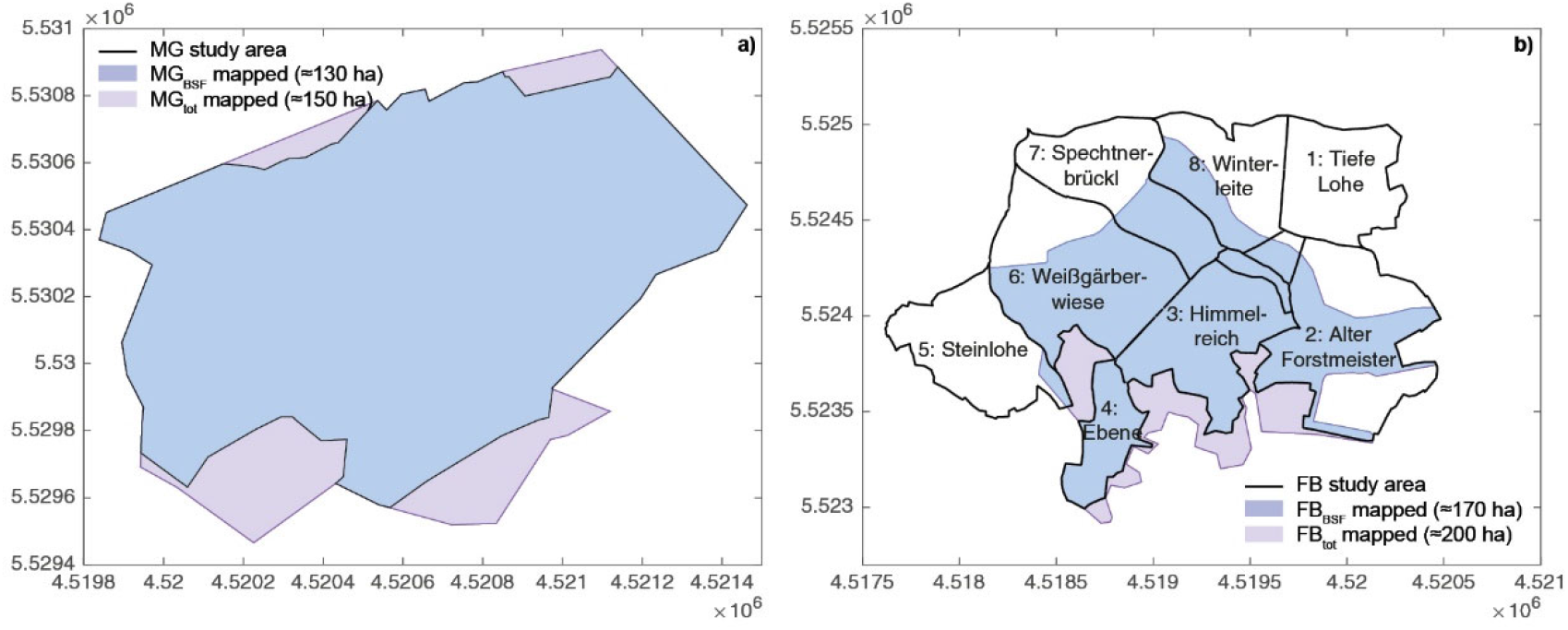
In total mapped areas (MG_tot_; FB_tot_) and mapped areas within the borders of BSF (MG_BSF_; FB_BSF_) for a) MG and b) FB study area.

### 2.3 Definition of variable and quasi-invariant site factors

To investigate “GeoBio-Interactions” in this study, we defined seven variable and four quasi-invariant factors. Variable factors are defined as factors that are influenced in a short time frame, such as physical nest parameters (e.g., NH and ND), nest building material, spatial distribution of RWA nests, RWA nest densities, or by human activities such as primary (TS_prime_) and secondary (TS_sec_) tree species, and tree age classes. Forest information was taken from the 10-year forest inventory and management plan provided by the Bayerische Staatsforsten AöR, Regensburg (BSF 2009) In addition, information on the location of RWA nests, e.g., within the forest stand, and herb layer at each RWA nest was collected and analyzed in the field.

Quasi-invariant factors are defined as long-lived factors that exist on geological time scales (some tens of thousands to some hundred million years). These factors are related to tectonic processes and are not influenced by human activities. In this study four main factors were selected: geochemical composition of the bedrock, geomorphology (terrain slope), exposure and tectonics. Information on these factors was collected and compiled from various sources, such as published geologic and tectonic maps (see Section “List of geologic maps” includes all authors of geologic maps that are indirectly cited by this term), and from the LMU, Munich earthquake catalogue (LMU 2020). For the analysis of the geomorphology, a digital terrain model (DTM; 1 m resolution) was used (LDBV 2008/2009).

### 2.4 Data analysis

Analyses were done with MATLAB R2018b (www.mathworks.com) and a geographic information system QGis (Version 3.10.7). Density plots of RWA nests were created using the code developed by Changyong (2020). A one-way analysis of variance (ANOVA) was applied to investigate variable and quasi-invariant factors in both study areas. Levene’s test was applied to investigate the similarity of variances between the parameters. Expected nest values were calculated proportional to the investigated forest area. We applied point distribution statistics to investigate whether RWA nests were evenly or randomly distributed or clustered by applying X^2^–test. Terrain slope was calculated by applying the slope algorithm of the processing-raster terrain analysis (QGIS 3.10.7) on the 1-m DTM (LDBV 2008/2009). Two slope classifications were determined: a) low slope ≤ 5° (gradient: 0–8.7 %) and b) steep slope ≥ 5° (gradient ≥ 8.7 %). Exposures were calculated using the aspect algorithm of the processing-raster terrain analysis (QGIS 3.10.7).

## 3 Results

### 3.1 Variable factors

Information on primary (TS_prime_), secondary tree species (TS_sec_) and tree ages were only available for the BSF (BSF 2009) but not for areas of other forest owners (OFO), e.g. privately owned or municipal forests. Therefore, the variable factors are discussed only for MG_BSF_ and FB_BSF_. In accordance with BSF, the medium tree age class was chosen for all analyses.

#### 3.1.1 Physical nest parameters

In total, 5,157 (FB: 2,829) and (MG: 2,328) RWA nests (*Formica polyctena*) were mapped in both study areas, of which ≈86 % (FB_BSF_ 2,425) and ≈92 % (MG_BSF_ 2,143) had been mapped in BSF forests; 1.3 % (MG_BSF_) and 8.3 % (FB_BSF_) were abandoned. The remaining data (FB_OFO_ ≈14 %; MG_OFO_ ≈8 %) can be attributed to OFO. Most nests (MG_BSF_ ≈66 %; FB_BSF_ ≈74 %) were start-up and short nests (0.01–0.5 m height), and one fifth (FB_BSF_ ≈22 %) and one third (MG_BSF_ ≈27 %) were medium sized nests (0.51–1.0 m). The tallest nests (≥1.01 m) were located outside the BSF forests (MG_OFO_ ≈16 %; FB_OFO_ ≈5 %). The largest nest diameters (≥1.51 m) were mapped in FB_BSF_ forest section No. 4 “Ebene” (≈14 %), No. 3 “Himmelreich” (≈11 %), and in MG_OFO_ (≈22 %; Tab. 2). Up to three quarters (MG_BSF_ ≈72 %; FB_BSF_ ≈53 %) of all mapped nests were located within forest stands and natural regeneration areas, and up to one third (FB_BSF_ ≈31 %; MG_BSF_ ≈22 %) at forest roads and skid trails. Open space areas such as clearings within the forest, meadows, fields, ponds or islands were not preferred by RWA nests (MG_BSF_ ≈7 %; FB_BSF_: 16 %). Blue berries [*Vaccinium myrtillus*] were the main herbs (MG_BSF_ ≈69 %; FB_BSF_ ≈68 %) accompanying start-ups, short and medium sized nests, followed by moss [*Bryophta*; MG_BSF_ ≈21 %; FB_BSF_ ≈36 %]. Gramineous plants [*Poales*] were also observed (MG_BSF_ ≈2 %; FB_BSF_ ≈31 %). Other typical plants of the herb layer, such as ferns, clover, lupines or nettles only played a minor role.

**Table 2.** Descriptive statistics of mapped nest height (NH) and diameter (ND) classes in a) MG_BSF_ and b) FB_BSF_ and forest areas of other forest owners (OFO). Additionally, data for each investigated FB forest sections are listed. –.= not available

| **a) RWA nest height (NH) classes [m]** |  | **0.01–0.10** | **0.11–0.50** | **0.51–1.00** | **1.01–1.50** | **1.51–2.00** | **–** |
| --- | --- | --- | --- | --- | --- | --- | --- |
| **MG (mapped area)** | **n** | **(%)** | **(%)** | **(%)** | **(%)** | **(%)** | **(%)** |
| $MG_{BSF}$ ($\approx 130$ ha) | 2,143 | 12.6 | 53.2 | 27.3 | 6.3 | 0.6 | – |
| $MG_{OFO}$ ($\approx 20$ ha) | 185 | 5.9 | 53.5 | 24.9 | 10.8 | 4.9 | – |
| $MG_{tot}$ ($\approx 150$ ha) | 2,328 | 12.0 | 53.3 | 27.1 | 6.7 | 0.9 | – |
| **FB (mapped area)** |  |  |  |  |  |  |  |
| $FB_{BSF}$ ($\approx 170$ ha) | 2,425 | 16.3 | 57.6 | 21.8 | 4.2 | 0.0 | – |
| $FB_{OFO}$ ($\approx 30$ ha) | 404 | 15.8 | 49.0 | 25.5 | 9.2 | 0.5 | – |
| $FB_{tot}$ ($\approx 200$ ha) | 2,829 | 16.3 | 56.4 | 22.3 | 4.9 | 0.1 | – |
| 2: Alter Forstmeister ($\approx 31$ ha) | 233 | 13.3 | 56.2 | 24.9 | 5.6 | – | – |
| 3: Himmelreich (42 ha) | 736 | 15.4 | 60.9 | 20.0 | 3.8 | – | – |
| 4: Ebene (17 ha) | 401 | 11.0 | 52.4 | 31.9 | 4.5 | 0.2 | – |
| 5: Steinlohe (2 ha) | 12 | 8.3 | 41.7 | 50.0 |  | – | – |
| 6: Weißgärberwiese (44 ha) | 617 | 20.1 | 59.8 | 16.5 | 3.6 | – | – |
| 7: Spechtnerbrückl (14 ha) | 282 | 19.5 | 53.5 | 22.3 | 4.6 | – | – |
| 8: Winterleite (17 ha) | 144 | 19.4 | 57.6 | 17.4 | 5.6 | – | – |
| **b) RWA nest diameter (ND) classes [m]** |  |  |  |  |  |  |  |
|  |  | **0.01–0.50** | **0.51–1.00** | **1.01–1.50** | **1.51–2.00** | **2.01–2.50** | **>2.51** |
| **MG (mapped area)** | **n** | **(%)** | **(%)** | **(%)** | **(%)** | **(%)** | **(%)** |
| $MG_{BSF}$ ($\approx 130$ ha) | 2143 | 41.6 | 30.3 | 17.5 | 7.5 | 1.0 | 2.1 |
| $MG_{OFO}$ ($\approx 20$ ha) | 185 | 33.0 | 30.3 | 14.6 | 9.7 | 4.3 | 8.1 |
| $MG_{tot}$ ($\approx 150$ ha) | 2328 | 40.9 | 30.3 | 17.2 | 7.7 | 1.2 | 2.6 |
| **FB (mapped area)** |  |  |  |  |  |  |  |
| $FB_{BSF}$ ($\approx 170$ ha) | 2425 | 44.2 | 28.6 | 18.4 | 5.4 | 1.7 | 1.6 |
| $FB_{OFO}$ ($\approx 30$ ha) | 404 | 39.6 | 23.8 | 23.3 | 8.4 | 2.7 | 2.2 |
| $FB_{tot}$ ($\approx 200$ ha) | 2829 | 43.5 | 27.9 | 19.1 | 5.9 | 1.9 | 1.7 |
| 2: Alter Forstmeister ($\approx 31$ ha) | 233 | 46.4 | 32.6 | 19.7 | 1.3 | 0.0 | 0.0 |
| 3: Himmelreich (42 ha) | 736 | 41.3 | 29.1 | 18.5 | 6.4 | 2.9 | 1.9 |
| 4: Ebene (17 ha) | 401 | 33.7 | 27.7 | 24.9 | 8.7 | 1.7 | 3.2 |
| 5: Steinlohe (2 ha) | 12 | 50.0 | 25.0 | 25.0 | 0.0 | 0.0 | 0.0 |
| 6: Weißgärberwiese (44 ha) | 617 | 53.8 | 27.4 | 12.6 | 4.7 | 1.0 | 0.5 |
| 7: Spechtnerbrückl (14 ha) | 282 | 39.4 | 28.4 | 22.3 | 5.0 | 2.5 | 2.5 |
| 8: Winterleite (17 ha) | 144 | 52.8 | 27.8 | 14.6 | 2.8 | 0.7 | 1.4 |

#### 3.1.2 Primary and secondary tree species, tree age

Nest height and diameter were significantly influenced by forest type, which consisted of different primary (TS_prime_) and secondary tree species (TS_sec_), age classes, nest building material and the geochemical composition of bedrock, but not by geomorphology as confirmed by the results of one-way ANOVA (Tab. 3). Levene’s test confirmed the similarity of variances between the parameters studied.

**Table 3.** Results of one-way ANOVA for physical nest parameters (diameter and height) for variable factors for a) MG_BSF_ and b) FB_BSF_ and quasi-invariant factors for c) MG_tot_ and d) FB_tot_. Parameters that significantly differ are set in bold. – = not available.

|  | *F* | df1 | *p* |  | *F* | df1 | *p* |
| --- | --- | --- | --- | --- | --- | --- | --- |
| **a) MGBSF** (No. of RWA nests = 2143) |  |  |  | **b) FBBSF** (No. of RWA nests = 2425) |  |  |  |
| **Variable factors** |  |  |  | **Variable factors** |  |  |  |
| **Forest type (with regard to TS<sub>prime</sub>)** |  |  |  | **Forest type (with regard to TS<sub>prime</sub>)** |  |  |  |
| Height | 3.206 | 5 | **0.007** | Height | 2.850 | 3 | **0.036** |
| Diameter | 2.234 | 5 | **0.049** | Diameter | 2.646 | 3 | **0.048** |
| **Forest type (with regard to TS<sub>sec</sub>)** |  |  |  | **Forest type (with regard to TS<sub>sec</sub>)** |  |  |  |
| Height | 1.960 | 3 | 0.118 | Height | 5.424 | 5 | **<0.001** |
| Diameter | 2.315 | 3 | 0.074 | Diameter | 3.108 | 5 | **0.008** |
| **Age class** |  |  |  | **Age class** |  |  |  |
| Height | 3.665 | 6 | **0.001** | Height | 6.285 | 6 | **<0.001** |
| Diameter | 8.929 | 6 | **<0.001** | Diameter | 10.545 | 6 | **<0.001** |
| **Nest building material** |  |  |  | **Nest building material** |  |  |  |
| Height | 23.983 | 2 | **<0.001** | Height | 5.627 | 2 | **0.004** |
| Diameter | 64.164 | 2 | **0.000** | Diameter | 15.953 | 2 | **<0.001** |
| **a) MG<sub>tot</sub>** (No. of RWA nests = 2328) |  |  |  | **b) FB<sub>tot</sub>** (No. of RWA nests = 2829) |  |  |  |
| **Quasi-invariant factors** |  |  |  | **Quasi-invariant factors** |  |  |  |
| **Geochemical composition of bedrock** |  |  |  | **Geochemical composition of bedrock** |  |  |  |
| Height | 3.594 | 3 | **0.013** | Height | 5.130 | 1 | **0.024** |
| Diameter | 4.949 | 3 | **0.002** | Diameter | 19.879 | 1 | **<0.001** |
| **Geomorphology** |  |  |  | **Geomorphology** |  |  |  |
| Height | 0.015 | 3 | 0.997 | Height | 1.501 | 3 | 0.212 |
| Diameter | 1.288 | 3 | 0.278 | Diameter | 0.487 | 3 | 0.691 |

Mature (≥80–140 years) pine (TS_prime_)-dominated forests are the preferred location for RWA nests (MG_BSF_ ≈70 %; FB_BSF_ ≈60 %). Here, 3 times (FB_BSF_) and 7 times (MG_BSF_) more RWA nests were mapped and a fifth of start-ups to short nests with small diameters (up to 0.5 m) compared to mature spruce (TS_prime_)-dominated forests (≥ 80 years; only ≈6 % of RWA nests). In spruce-dominated forests, the preferred tree age classes were 20, 40, 60 and 120 years. Here, the number of RWA nests was 2.3 times (MG_BSF_) and 1.3 times (FB_BSF_) lower than in pine-dominated forests. Deciduous trees, e.g. beech, hornbeam, white birch, were no relevant TS_prime_ due to low RWA nest numbers (Fig. 4). The percentages of nest diameter (ND) classes in mature forests (≥ 80 years) were very similar to pine and spruce (TS_prime_) in both study areas: small nests ≈45 %; medium nests≈28 %, and large–very large nests ≈27 %.

**Fig 4.**
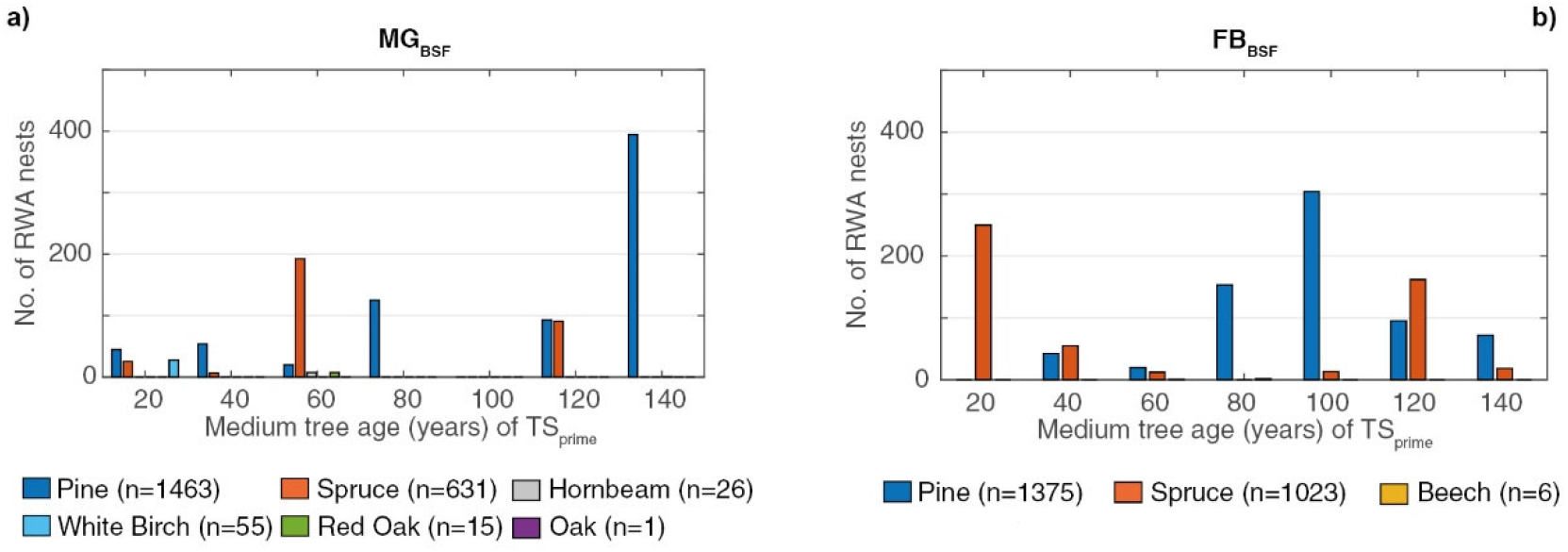
Numbers of RWA nests versus medium tree age of primary tree species (TS_prime_) for a) MG_BSF_ and b) FB_BSF_

A comparison of TS_prime_ and TS_sec_ with RWA nest numbers confirmed our results that RWA preferred pine-dominated forests. Also, the RWA nest numbers of all height classes are almost twice as high in pine-dominated mixed coniferous forests (pine=TS_prime_/spruce=TS_sec_) as in an inverse situation (spruce = TS_prime_/pine = TS_sec_). In addition, the majority of short to medium sized nests (MG_BSF_ ≈83 %; FB_BSF_ ≈80 %) are found in pine-dominated mixed coniferous forests. Other tree species like oak, red oak, beech or Douglas fir do not play a decisive role.

#### 3.1.3 Nest building material

The preferred nest building material in all nest height classes were fresh to slightly decomposed pine needles in pine-dominated mixed coniferous forests (pine=TS_prime_/spruce=TS_sec_). Even in spruce-dominated forest, pine needles-dominated (FB_BSF_). In MG_BSF_, RWA nests showed a slightly higher preference for spruce needles (≈44 %) compared to pine needles (≈35 %; Fig. 5).

**Fig 5.**
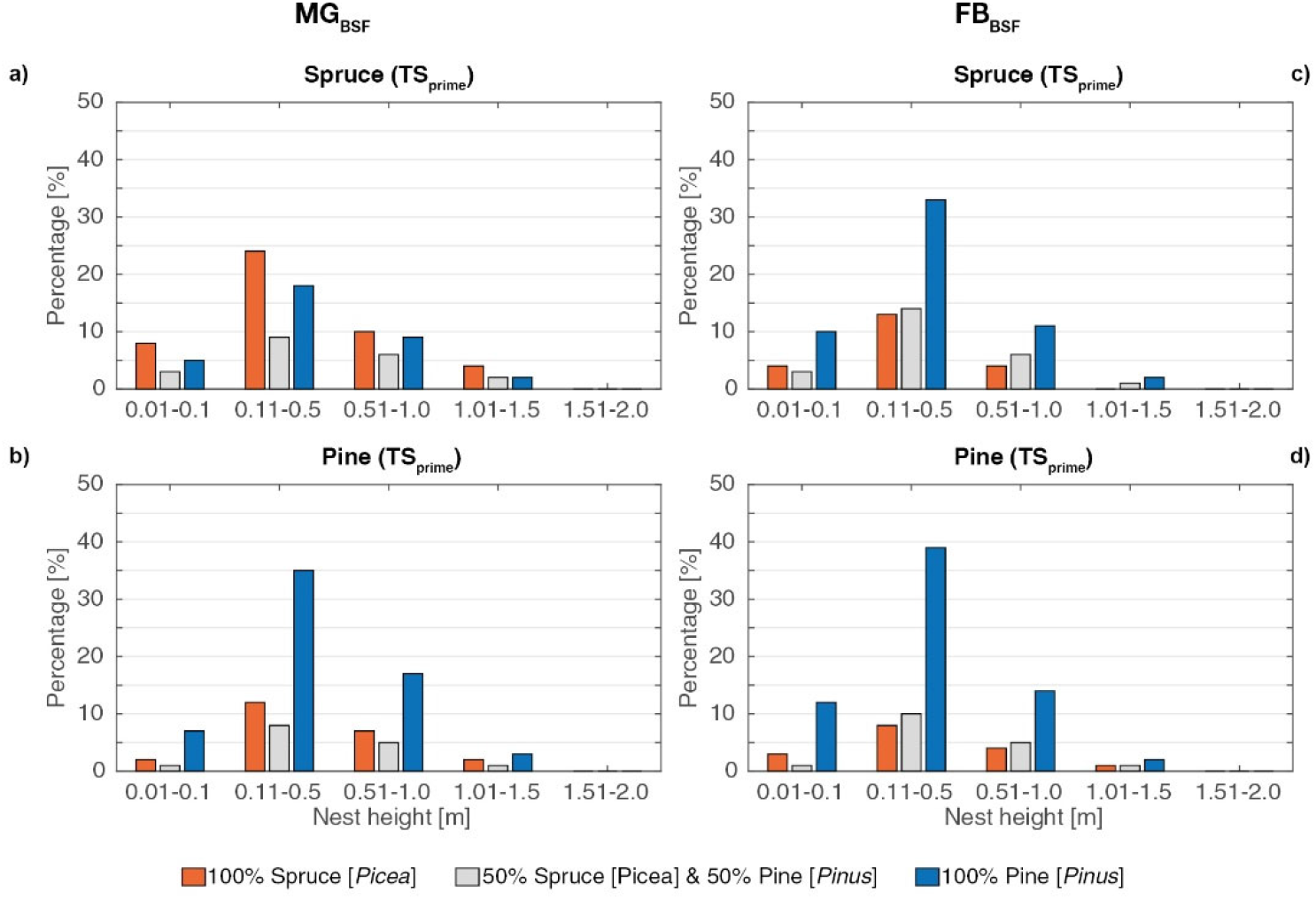
Nest height classes[m] versus type of nest material (%) for a,c) spruce, and b,d) pine as primary tree species (TS_prime_) in MG_BSF_ and FB_BSF._

#### 3.1.4 Spatial distribution and nest densities

In both study areas, RWA nests were spatially clustered, indicated by a nearest neighbor ratio <1 (MG_BSF_: 0.5; FB_BSF_ 0.3) and Z-statistic < -1.96 (Z_FB_: -82.84; Z_MG_: -71.64) at a significance level of 95 %. In FB_BSF_, 10 % more nests were mapped than indicated by the expected number of RWA nests. Nest densities were very high: 16.5 nests/ha in MG_BSF_ and 14.3 nests/ha in FB_BSF_ (Tab. 4). The highest nest densities were found in FB_BSF_ forest sections No. 4 “Ebene” (≈24 nests/ha) and No. 7 “Spechtnerbrückl” (20 nests/ha). Mature (≥ 80 years) pine-dominated mixed coniferous forests (pine= TS_prime_/spruce= TS_sec_) showed median nest densities of 13.3 nests/ha (MG_BSF_) and 11.9 nests/ha (FB_BSF_), which are comparable to those for spruce-dominated mixed coniferous forests (spruce=TS_prime_/pine=TS_sec_; MG_BSF_ 10.9 nests/ha; FB_BSF_ 13.3 nests/ha). Furthermore, a striking radial nest distribution pattern (≈440 m in diameter) was observed in the NE of the FB_BSF_ forest section No. 3 “Himmelreich”.

**Table 4.**
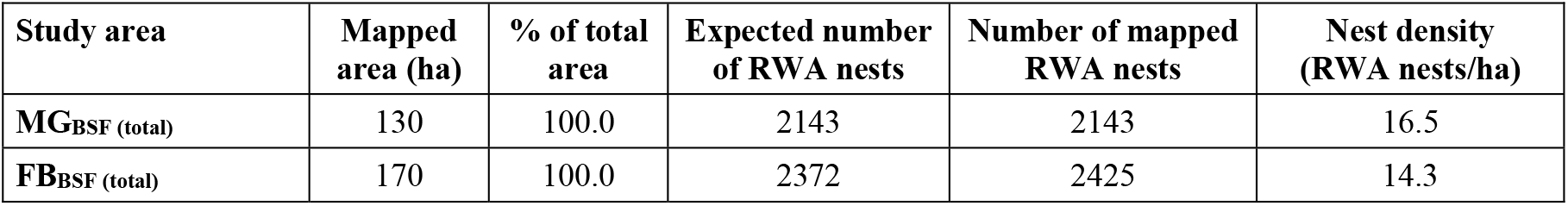
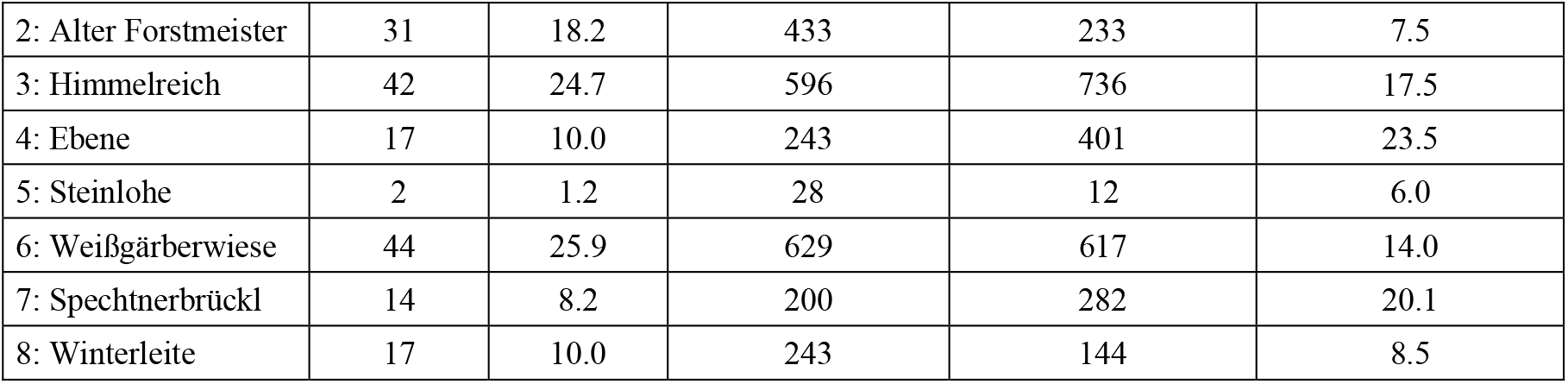
Descriptive statistics of mapped forest areas, percentage of total mapped area, expected numbers of RWA nests, numbers of mapped nests in the field, density of RWA nests (nests/ha) for MG_BSF_ and FB_BSF_

| Study area | Mapped area (ha) | % of total area | Expected number of RWA nests | Number of mapped RWA nests | Nest density (RWA nests/ha) |
| --- | --- | --- | --- | --- | --- |
| MG<sub>BSF</sub> (total) | 130 | 100.0 | 2143 | 2143 | 16.5 |
| FB<sub>BSF</sub> (total) | 170 | 100.0 | 2372 | 2425 | 14.3 |
| 2: Alter Forstmeister | 31 | 18.2 | 433 | 233 | 7.5 |
| 3: Himmelreich | 42 | 24.7 | 596 | 736 | 17.5 |
| 4: Ebene | 17 | 10.0 | 243 | 401 | 23.5 |
| 5: Steinlohe | 2 | 1.2 | 28 | 12 | 6.0 |
| 6: Weißgärberwiese | 44 | 25.9 | 629 | 617 | 14.0 |
| 7: Spechtnerbrückl | 14 | 8.2 | 200 | 282 | 20.1 |
| 8: Winterleite | 17 | 10.0 | 243 | 144 | 8.5 |

### 3.2 Quasi-invariant factors

Analyses of quasi-invariant factors based on the field survey and included all data on RWA nests for BSF and OFO forest areas (henceforth MG_tot_ and FB_tot_), as these factors are independent of the BSF data base.

#### 3.2.1 Geochemical composition of bedrock

In total, 2,328 (MG_tot_) and 2,829 (FB_tot_) RWA nest were mapped (Tab. 5). The geochemical composition of the bedrock had a significant influence on nest heights and diameters in both study areas (Tab. 3 and 5). In MG, the Mitterteicher/Steinwald Granite (≈13 %) and the Miocene and Miocene-Pliocene (≈12 %) sedimentary cover showed comparable percentage of nest start-ups. For all other nest height and diameter classes there were no major differences. Due to the low RWA nest numbers (18) at the Pliocene sedimentary lens, the statistics cannot be compared to the other geological units (Tab. 5). In FB, ≈12 % RWA nests were mapped on the younger Mitterteich/Steinwald Granite (≈15 ha; Muscovite-Monzogranite) and 2,496 (≈88 %) RWA nests on the older Falkenberger Granite (≈184 ha; Andalusite-Sillimanite-Monzogranite) in total. The Falkenberger Granite had slightly more nest start-ups (≈17 %) compared to the Mitterteich/Steinwald Granite. Tall to very tall nests (≥ 1.01 m) and RWA nests with large diameters (≥ 1.51 m) were more frequent at the Mitterteicher/Steinwald Granite (Muscovite-Monzogranite) compared to the older Falkenberg Granite (Andalusite-Sillimanite-Monzogranite; Tab. 5).

**Table 5.**
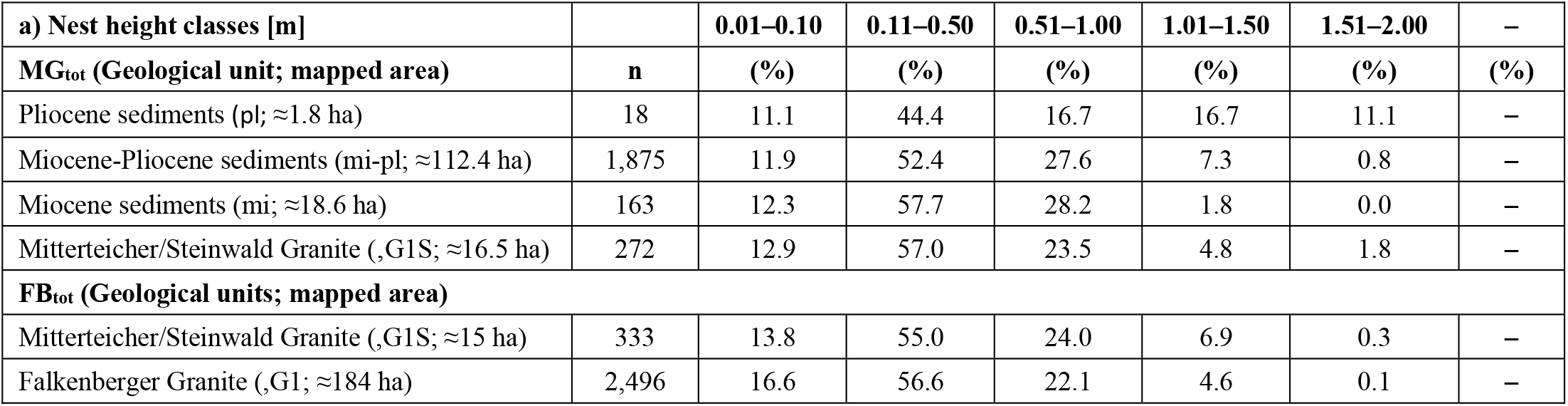
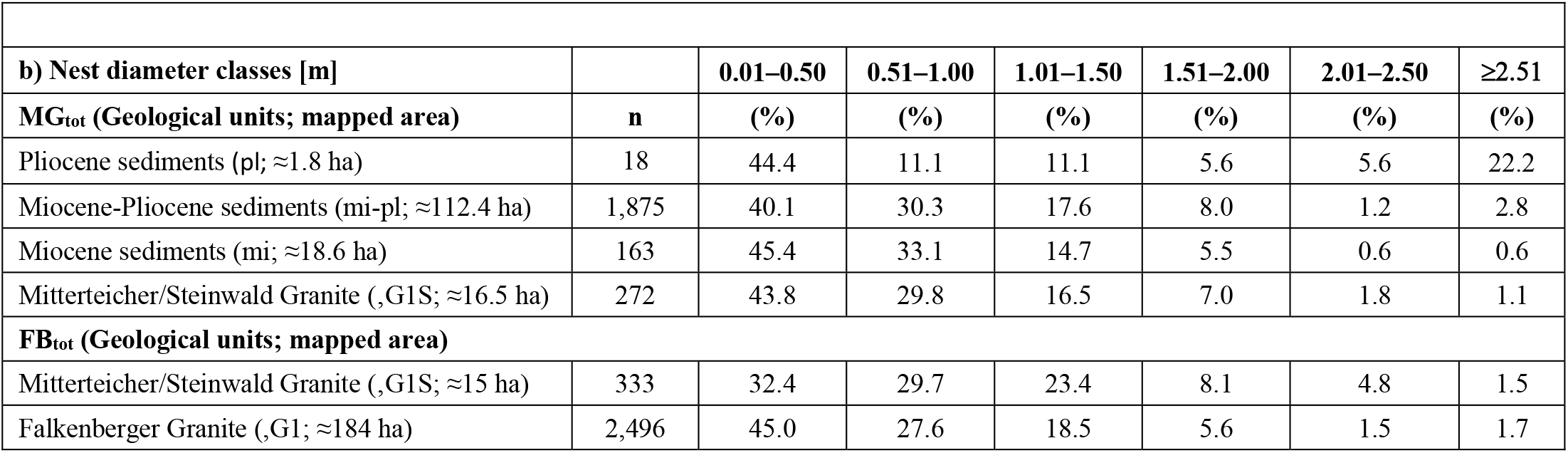
Comparison in percentage [%] for a) nest height and b) diameter classes for different geological units in MG_tot_ and FB_tot_. – = not available.

| a) Nest height classes [m] |  | 0.01–0.10 | 0.11–0.50 | 0.51–1.00 | 1.01–1.50 | 1.51–2.00 | – |
| --- | --- | --- | --- | --- | --- | --- | --- |
| MG<sub>tot</sub> (Geological unit; mapped area) | n | (%) | (%) | (%) | (%) | (%) | (%) |
| Pliocene sediments (pl; $\approx 1.8$ ha) | 18 | 11.1 | 44.4 | 16.7 | 16.7 | 11.1 | – |
| Miocene-Pliocene sediments (mi-pl; $\approx 112.4$ ha) | 1,875 | 11.9 | 52.4 | 27.6 | 7.3 | 0.8 | – |
| Miocene sediments (mi; $\approx 18.6$ ha) | 163 | 12.3 | 57.7 | 28.2 | 1.8 | 0.0 | – |
| Mitterteicher/Steinwald Granite (,G1S; $\approx 16.5$ ha) | 272 | 12.9 | 57.0 | 23.5 | 4.8 | 1.8 | – |
| FB<sub>tot</sub> (Geological units; mapped area) |  |  |  |  |  |  |  |
| Mitterteicher/Steinwald Granite (,G1S; $\approx 15$ ha) | 333 | 13.8 | 55.0 | 24.0 | 6.9 | 0.3 | – |
| Falkenberger Granite (,G1; $\approx 184$ ha) | 2,496 | 16.6 | 56.6 | 22.1 | 4.6 | 0.1 | – |

| **b) Nest diameter classes [m]** |  | **0.01–0.50** | **0.51–1.00** | **1.01–1.50** | **1.51–2.00** | **2.01–2.50** | **≥2.51** |
| --- | --- | --- | --- | --- | --- | --- | --- |
| **MG<sub>tot</sub> (Geological units; mapped area)** | **n** | **(%)** | **(%)** | **(%)** | **(%)** | **(%)** | **(%)** |
| Pliocene sediments (pl; ≈1.8 ha) | 18 | 44.4 | 11.1 | 11.1 | 5.6 | 5.6 | 22.2 |
| Miocene-Pliocene sediments (mi-pl; ≈112.4 ha) | 1,875 | 40.1 | 30.3 | 17.6 | 8.0 | 1.2 | 2.8 |
| Miocene sediments (mi; ≈18.6 ha) | 163 | 45.4 | 33.1 | 14.7 | 5.5 | 0.6 | 0.6 |
| Mitterteicher/Steinwald Granite (,G1S; ≈16.5 ha) | 272 | 43.8 | 29.8 | 16.5 | 7.0 | 1.8 | 1.1 |
| **FB<sub>tot</sub> (Geological units; mapped area)** |  |  |  |  |  |  |  |
| Mitterteicher/Steinwald Granite (,G1S; ≈15 ha) | 333 | 32.4 | 29.7 | 23.4 | 8.1 | 4.8 | 1.5 |
| Falkenberger Granite (,G1; ≈184 ha) | 2,496 | 45.0 | 27.6 | 18.5 | 5.6 | 1.5 | 1.7 |

#### 3.2.2 Geomorphology and terrain exposure of RWA nests

Most of the terrain of MG_tot_ is smooth, with only a few sections of the terrain having steep slopes up to 25 °. In MG_tot_, one third (≈33 %) of start-up and short nests with ND≤ 0.5 m and tall nests (height and diameter: ≥ 1.0 m; ≈27 %) were on low slopes (≤ 5°). Steep slopes (≥ 5°) were preferred only by 7 % of the nests. However, two short nests with ND≤ 0.5 m were mapped on steep slopes (25 °). The relationships are the same for both, spruce- and pine-dominated coniferous forests (MG_BSF_) for the above-mentioned nest classes.

The geomorphology of FB_tot_ is different. Apart from a few flat areas, the terrain can be described as rugged with the steepest slopes up to 33 °, where three short nests with ND≤ 0.5 m were mapped. In FB_tot_, one fifth (≈19 %) of all mapped start-up and short nests with ND≤ 0.5 m were on low slopes (≤ 5°); and one quarter (≈25 %) on steep slopes (≥ 5°).

In spruce-dominated forests, RWA preferred steeper slopes for start-up and short nests with ND≤ 0.5 m (FB_tot_ ≈33 %). In pine-dominated forests, no difference between low (≈30 %) and steep (≈29 %) slopes was observed for the same nest classes. Taller nests (height and diameter: ≥ 1.0 m) were five times more frequent on steep slopes (≈30 %) than on low slopes (≈6 %) in spruce-dominated coniferous forests, but showed no differences for pine-dominated coniferous forests (≈20 % at slopes ≤ 5° and ≈17 % at slopes ≥ 5°).

In both study areas, exposure is not a relevant factor for RWA nests. In MG_tot_, the preferred directions are SE (≈7.4 % of RWA nests), SSE (7.0 % of RWA nests) and NNW (≈7.2 % of RWA nests); in FB_tot,_ the preferred directions were S (≈7.6 % of RWA nests), SSW (≈7.5 % of RWA nests), NNE (≈7.1 % of RWA nests), and N (≈7.0 % of RWA nests) directions were preferred. All other exposures were distributed almost equally among RWA nests in both study areas (MG_tot_: 5.0 %– 6.8 %; FB_tot_: 5.2–6.5 %; Fig. 6).

**Fig 6.**
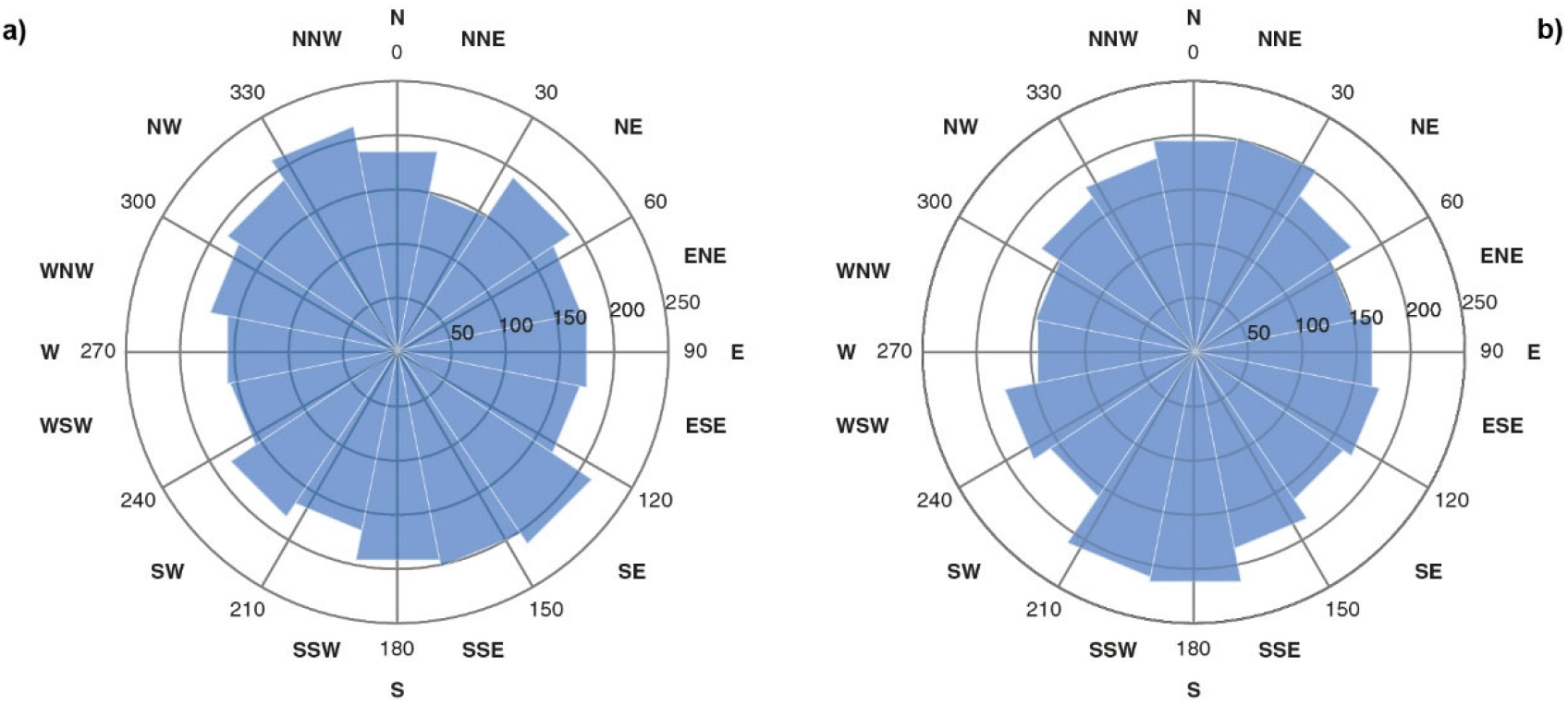
Preferred terrain exposure of RWA nests in a) MG_tot_ (n=2,328) and b) FB_tot_ (n=2,829) study area

#### 3.2.3 Tectonics

Density plots calculated for both study areas (Changyong 2020) showed a mostly NW-SE spatial distribution pattern of RWA nests, paralleling the present-day main stress direction in the area. Additionally, WNW-ESE directions were found in the western part of MG study area (Fig. 7). During the field campaigns, no earthquakes occurred in both study areas.

**Fig 7.**
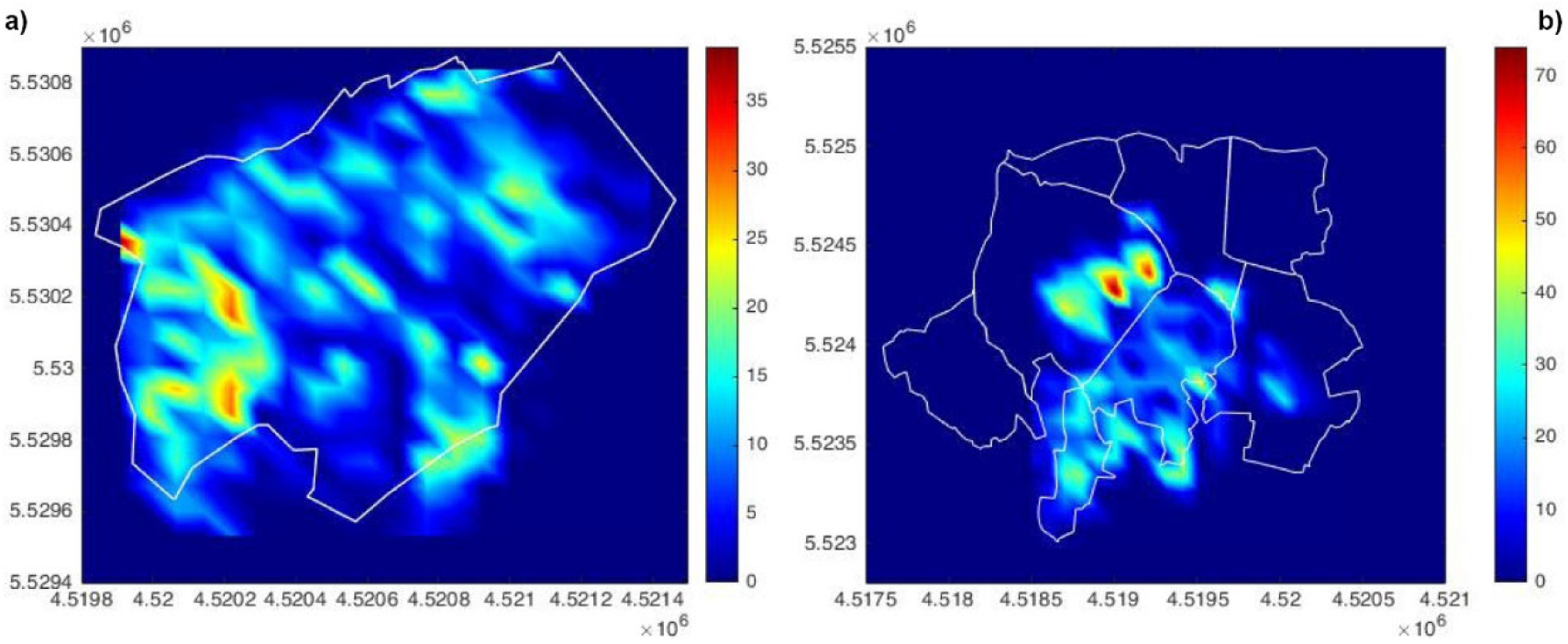
Density plots of RWA nests in a) MG and b) FB study area.

## 4 Discussion

Sufficient information on occurrence and spatial distribution is required to protect RWA as keystone species, ecosystem engineers, and biocontrol agents in an integrally managed forest. The area-wide mapping carried out in this study based on presence/absence data provided for the first time a reliable, GPS-based RWA nests database for both study areas. Since some RWA are considered species of conservation concern (e.g., BfN 2012; IUCN 2015), it is necessary to further apply effective conservation measures for RWA species in forest management. So far, preparatory work of the BSF for logging activities includes labeling of tree trunks near a RWA nest in 1.5 m height with neon colors, that can be recognized by forest workers from all directions. No logging activities or processing of trees are permitted in the vicinity of such labelled trees. With the recent GPS-based database, containing the area-wide distribution pattern of the RWA nests, the BSF can achieve its overall goal to further protecting the RWA nests. Future forest work will use this database to identify specific forest areas with RWA nests and nest clusters. In these specific areas, nature conservation will be given higher priority than the use of trees. Since the spatial distribution patterns of RWA nests may change over time, we suggest to re-map these areas within five years to quantify population changes in both study areas.

### 4.1 Variable factors

#### 4.1.1 Physical nest parameters, dominant tree species and tree age classes

Red wood ants of the *Formica rufa*-group are an ecologically indisputably successful species (Czechowski & Vesplänen 2009). In entomological studies, e.g., tree species and characteristics (Gibb et al. 2016) or type of forest management (Sorvari & Hakkarainen 2007) are regarded as decisive for RWA occurrences. Recent studies have found that stand age and the associated differences in forest structure have significant effects on abundance, nest size and behavior of ants (e.g., Punttila 1996; Gibb & Johansson 2010). According to Gibb & Johansson (2010), middle-aged forest stands (30–40 years old) in managed boreal forests showed the fewest and smallest ant nests and the lowest ant activity. Sondeij et al. (2018), reported the highest nest densities in 101–120 and 181–200-year-old forest stands and in forests younger than 20 years, suggesting young forests with an open canopy promote preferred habitats for nest settlements. In contrast, Vandegehuchte et al. (2017) reported that RWA mainly dependent on forest structure and conifer abundance. Either forest fragment size, distance to forest edges, nor shrub diversity were addressed as important factors.

Our findings showed, that the most favored tree age classes were mature pine-dominated forests (≥80–140 years) with 3–7 times more RWA nests compared to mature spruce-dominated forests (≥ 80 years). Nest numbers were 1.3–2.3 times lower in spruce-dominated forests with tree age classes of 20, 40, 60 and 120 years. This is in contrast to the findings by Sondeij et al. (2018), though mainly

*F. polyctena* (75 % of observed nests) was in the focus of this study and also in contrast to findings by Wiśniewski (1969), who observed lower nest densities in 60–100-years-old pine forests. Our findings are consistent with the life history of the polygynous *F. polyctena* which are more abundant in mature, closed canopy forests (Pisarski & Czechowski 1994; Punttila 1996). Furthermore, we cannot confirm findings by Sondeij et al. (2018) and Punttila (1996), that smaller nest sizes, which could indicate younger nests are typical for young forests, and that an open canopy cover is favorable for colony establishment. We observed one-fifth (≈20 %) of start-ups to short nests with small diameters (up to 0.5 m) in pine-dominated mature forest (≥ 80 years). The age class might have an influence on the light conditions, which in turn affects the nests size (Sondeij et al. 2018), but then no tall–very tall nests should be observed in these age classes. Our findings show up to 9 % of these tall–very tall nests were found in pine-dominated mature forest (≥ 80 years). We also cannot confirm findings by Domisch et al. (2005) who did not observe RWA nests in 20-years old Scots pine stands. In our two study areas, 4 % of all nests were mapped in young (up to 20 years old) pine forests, although the forest was managed in both study areas. One reason for this could be that in such forests, if necessary, a small number of trees is logged to promote light on the forest floor and to create sunny places for, e.g. reptiles or amphibians. Probably RWA might also benefit from of this conservation measures. It is further suggested that previous findings in entomological studies could also be a result of mapping approach, e.g. transects, plots. From our experience we suggest that only area-wide mapping and presence/absence data reveal the true results of RWA nests in forest habitats and provide an applicable database for further improved knowledge of ant habitats.

#### 4.1.2 Nest building material

RWA ants collect and concentrate organic material from the forest floor for the construction of their mostly conical above-ground nests (Frouz et al. 2005; Domisch et al. 2007). Conifer needles (Gösswald 1989), small branches and other plant material as leaf-litter and other detritus which can affect the nutritional and physical properties of the soil in a variety of ways (e.g., Frouz et al. 2005; Domisch et al. 2007) have been reported as favorable nest building material.

Our results showed that the preferred nest building material in both study areas and for almost all nest heights was pine needles. Even in spruce-dominated forests RWA tried to build their nest with longer pine needles. We observed pine needles on all nest forms, steeply conical, but also on flat forms. Such a clear preference for pine needles as nest building material, even though pines are rare in a forest stand, has not been observed so far. There could be mechanical and chemical reasons for the preference of pine needles: Pine needles may have a mechanical advantage in size and structure over individual spruce needles, as pine needles are usually found as a bundle of 2–3 extremely narrow needles per sheath covering the base of the needle bundle. In addition, a greater thickness of outer epidermal walls of pine needles can support mechanical strength and protect the pine needles from desiccation, especially in winter (Jankowski et al. 2017). This can support the durability of the nest building material. In addition, it is suggested that start-ups and short nest could have a more solid outer structure by building them from longer double or triple pine needles that are intertwined with each other compared to short loose spruce needles. Such a construction could also be more solid against wind driven processes. Furthermore, long intertwined pine needles could also strengthen steep conical nests and contribute to the stability of the nest. During the chemical decomposition of solid organic components, cellulose dominate in pine needle litter, which has a static function in plants as tear-resistant fibers, while lignin dominates in spruce (Johansson 1995). This could additionally contribute to longevity of the nest cover. From a chemical point of view, the thick-walled epidermis contributes to the defense against herbivores by forming specialized resin channels (Jankowski et al. 2017). Resin is an important component of RWA nests because of its stabilizing and antibacterial effect (Christe et al. 2003). Our results also support findings by Gibb et al (2016), that ants (*F. aquilonia*) harvested fresh needles directly from pine or spruce trees, especially in in older forests stands, which indicates better needle quality and lower activity of microorganisms on fresh needles.

#### 4.1.3 Nest densities

Things as conspicuous as ant nests can be easily overlooked. The findings on nest densities in Eurasian forests are highly variable and range from 0.02–0.76 (Sondeij et al. 2018) to 5 nests/ha (Risch et al. 2016). A recent study showed that the detection probability of red wood ant nests is surprisingly variable and depends on a) observer experience, b) nest size, c) mapping procedure (complete inventory with presence/absence data, random sampling, systematic versus opportunistic procedure), and d) area size, but not on habitat characteristics (forest type, local vegetation; Berberich et al. 2016c). Observer experience is crucial to detect even start-up and small nests RWA nests. A possible explanation for the large differences among studies on nest densities could be that no information was given on the observer(s) experience and on the estimation for detection probability. Very experienced observers like the first and second authors, who mapped more than 25,000 RWA nests in selected areas of Germany (e.g., Black Forest, Eifel area), Switzerland, Austria and the United States of America, found twice as many short RWA nests, 33% more tall ones and 66% more with smaller diameters (up to 50 cm) than beginners (Berberich et al. 2016c). Due to this demonstrably high detection probabilities of both observers (G.M.B. and M.B.B.; Berberich et al. 2016c) the high numbers of RWA nests in both study areas can be explained. Furthermore, the detection of RWA nest depends on the mapping procedure independent from the size of the area to be investigated. An area-wide complete inventory, as carried out in this study, always leads to more accurate results in contrast to transects, especially at distances of 20–50 m (e.g., Sondeij et al. 2018), for start-ups and short nests are easily overlooked. Additionally, recent studies (Berberich et al. 2016a, 2019) have shown, that RWA nests are not evenly distributed in the field, but are clumped and related to tectonically active fault zones. The hypothesis of an even spatial distribution of RWA nests and a coordinated mapping approach based on this leads to inaccurate data. Furthermore, very small areas with high numbers of RWA nests lead to very high, unrealistic nest densities compared to larger areas with lower nest numbers, as shown by Punttila & Kipilänen (2009). Such results are not comparable and lead to misinterpretations. We therefore suggest nest densities (nests/ha) are not suitable for regional comparison of RWA nest occurrences, as long as there is no comparable mapping approach, mapping is not carried out area-wide and no data on presence/absence have been collected.

### 4.2 Quasi-invariant factors

#### 4.2.1 Geochemical composition of bedrock and geomorphology

In the last century, entomologists reported that the spatial distribution of RWA nests depends on geological structures (Gösswald 1939; Rammoser 1961; Eichhorn 1962). Wellenstein (1929) found that RWA nests are not located at preferred sites within a forest stand. Recent studies investigating GeoBio-interactions showed that RWA nest occurrences are highly correlated with Radon degassing from rocks and soils into the atmosphere as part of the radioactive decay of the unstable isotopes of Uranium (U) (Berberich et al. 2016a, 2016b, 2018a, 2018b). In both study areas the U and Thorium (Th) concentrations of the granites indicate a very high natural Rn potential (MG: 51.8–177.2 kBq/m^3^; FB: 67.3–211.9 kBq/m^3^; BfS 2020). Soil profiles over weathered granites from the Upper Palatinate reflected the good adsorption properties of clay minerals and accumulation of U-bearing minerals, e.g. zirconium, monazite due to the strong increase in U-content in the grain fractions < 0.125 mm (Kemski et al. 2012). Our findings suggest that the geochemical composition of bedrock, in particular its overall high radioactive potential, might be the key factor in the distribution patterns for RWA nests. This hypothesis was already established in previous studies and could now be confirmed (Berberich et al. 2016a, 2016b, 2018a, 2018b). In both study areas, the small U-bearing soil horizons on the Variscan Granite but also the granites with their structurally determined pathways, both of which emit high Rn concentrations, are in direct contact with the RWA nests. In addition, temperature is an important factor influencing the biology, metabolic rate, growth, development, colony fitness and survival of poikilotherm RWA (Wilson 1971). Previous findings report that RWA can actively thermoregulate the nest due to accurate perception of temperature and evolution of temperature preferences. Besides insolation and heat exchange with air and soil, as well as endogenous heat generated by the ant colony, ants use metabolic heat as an additional source to maintain thermal homeostasis in the nest (Kadochová & Frouz 2014; Stukalyuk et al. 2020). Findings of a recent study by Stukalyuk et al. (2020) showed that heat loss of the nest depends on its size, the volume/nests ratio, and the condition and material of the cover layer. Furthermore, insolation does not cause the inner parts of the nests to heat up more than the temperature of the ground at the same depth. Therefore, *F. polyctena* nests with a diameter < 0.5 m are not capable of heating and are not autonomous at air temperatures up to +18 °C, nests diameters of 0.7 m can function normally at air temperature above +2 °C, nests with a nest diameter of > 0.8 m are able to maintain a constant optimum temperature in the nest (Stukalyuk et al. 2020). However, the question why start-up and short nests survive despite missing auto-thermoregulation in the winter, and further develop into tall nests and finally into polydomous colonies, is still unresolved. It is suggested that there is a process between RWA and the underlying bedrock material, especially in the Oberpfalz region, with short vegetation periods of < 140 days, 30 icy day/year and high precipitation rates (average 700–800 mm/year; LK-Tir 2020). The Late Variscan granites in both study areas have different petrogenesis and geochemical composition with regard to U and Th. The heat generated by the radioactive decay of the unstable isotopes U and Th, and potassium (K) is the largest internal heat source on Earth. In situ measurements of the Falkenberger Granite in 2015/2016 revealed U concentrations of 14.4 ppm ± 3.2 ppm, Th concentrations of 40.7 ppm± 4.8 ppm, and a Th/U ratio of 2.9± 0.6. The granites in the study area show a relatively high heat production of 6.7± 1.5 µW/m^3^ compared to other Variscan granites in Europe and can therefore be addressed as moderately heat producing granites (Scharfenberg & De Wall 2016). Findings by Kirchner (2007) showed that RWA are extremely sensitive to temperature. They can discriminate temperature differences of 0.25 K. It is suggested that RWA consider the continuous moderate heat production by the granite and its primarily U-bearing minerals in soils as favorable settlement conditions. During winter hibernation, the continuous heat emission from the granites and soils as an “additional heat source” can support the survival of the colony, especially start-up and short nests with diameters <0.5 m that are not able to warm up and thermoregulate themselves. This could also explain why the number of start-ups and short nests with diameters <0.5 m was very high (FB_tot_ ≈44 %; MG_tot_ ≈41 %) in both study areas.

The geomorphology is an additional factor that determines Rn concentrations. Lower terrain slopes promote higher values and show the least seasonal variations (Kemski et al. 2012). The results of the one-way ANOVA showed that the geomorphology is of little importance in both study areas. We attribute this to the fact that the overall high Rn potential provides optimal conditions and promotes the spatial distribution of RWA nests, even though the terrain might be rugged and flat sections are rare (FB study area). However, the geomorphology in combination with a cold climate and high precipitation rates, as it is observed in the Oberpfalz area, supports occurrence and distribution patterns of RWA nests, because rainy seasons (spring, autumn) lead to an increase in the radon concentration in the soil due to reduced migration. In winter, during snow cover, when the sealing effect is longer and more pronounced, and during frost periods, Rn emission into the atmosphere can be completely prevented, resulting in high radon concentrations in soil gas (Kemski et al. 2012). This is in contrast to previous studies, which suggested variable factors, such as canopy cover and edge (e.g., Risch et al. 2008), fragmentation (Punttila & Kilpelainen 2009), tree species, characteristics and age (e.g., Gibb et al. 2016) as relevant primary key factors. Our findings may also explain, why the RWA nest are found in extremely high numbers in the Oberpfalz study areas in contrast to findings of e.g., Sondeij et al. (2018) in the Białowieża Forest. Here, the moraine highlands are of different geochemical composition and do not offer such a high Rn potential, because their subsoil was formed during the retreat of the Warta Glaciation (Jaroszewicz et al. 2019).

#### 4.2.2 Exposure of RWA nests

Previous findings, e.g. (Wellenstein 1990) indicate that RWA prefer sites with S exposures. Findings by Harterbrodt (1990) regarding habitat requirements suggest that especially *F. polyctena* nests were more often mapped on terrains with W and N exposure. Our findings confirm these results. In both study areas NW, NNW, N, NNE and NE exposures had been frequent. Overall, however, there were only very small differences between the different exposures in both study areas. Therefore, we hypothesize that the geochemical composition of the bedrock is the key factor for the spatial distribution of RWA nests, and exposure is less important.

#### 4.2.3 Tectonics, distribution patterns and earthquakes

In entomological studies, the spatial distribution pattern of RWA nests and their local occurrence are related to a range of selective factors, e.g. foraging and food supply (e.g., Iakovlev et al. 2017), reproductive success (e.g., Rosengren et al. 1987), or founding of new nests or the colonization of new territories (e.g., Czechowski & Vepsäläinen 2009). Recent studies focusing on GeoBio-Interactions, e.g., in Germany and Romania, alternatively postulated spatial distribution patterns of RWA to be governed by degassing tectonic fault systems. The large-scale spatial distribution of RWA nests directly reflects significant tectonic components of the present-day stress field and its accompanying conjugated shear systems (Berberich et al. 2016a; Del Toro et al. 2017; Berberich et al. 2019). Furthermore, a double-blind study on Jutland (Denmark) showed that RWA nests were eight times more likely to be found within 60 m of known tectonic faults than were random points in the same region but without nests (Del Toro et al. 2017). The spatially clustered distribution patterns of the observed nests in both study areas suggests a strong interaction between RWA nests and their quasi-invariant environment. In particular, the spatial NW-SE distribution pattern of RWA nests found for both study areas corresponded to a) the directionality of the present-day stress field in NW-SE direction (Heidbach 2016), b) the NW-SE trending direction of the Erbendorfer Line, that delimits two regions with a different tectono-metamorphic history in the North of both study areas (Glaser et al. 2007), and c) the NW-SE striking intrusion fault line for the granites (Scharfenberg & De Wall 2016). The additionally identified WNW-ESE and W-E directions indicated by RWA nests are identical to the WNW-ESE and W-E trending quartz dike systems south of the FB study area (Fig. 2a). For there is no further information on the location of small-scale fault systems in MG and FB study areas due to the vegetation cover, the distribution of RWA nests complements and clarifies the tectonic regime identified in previous tectonic studies. In addition, the radial distribution pattern of RWA nests, which was observed in the northeast of the forest section No. 3 “Himmelreich”, could be attributed to “onion-like” joints in the Falkenberger Granite, that are comparable to those observed in the former quarry beneath the ruin Flossenbürg castle, approx. 14.5 km southeast of the FB study area (Glaser et al. 2007; Fig. 8).

**Fig 8.**
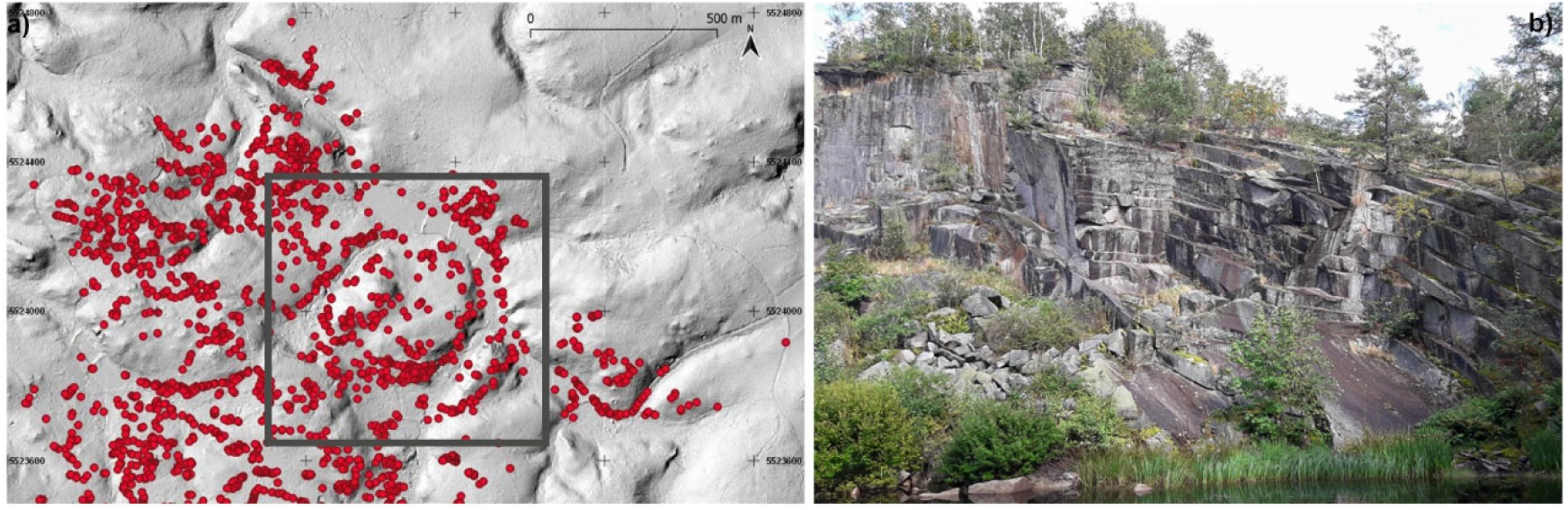
a) Spatial distribution of mapped RWA nests in FB_BSF_ based on the 1m DTM provided by the LDBV (2008/2009). Black square indicates a striking radial nest distribution pattern that can be explained by b) “onion-like” joints forming in the granite, as can be seen in the former quarry beneath the ruin Flossenbürg castle, approx. 14.5 km southeast of the FB Study area. Photo credit: M. Gibhardt.

The low number of weak to moderate earthquakes (LMU 2020) in the vicinity of both study areas does not play a decisive role (LMU 2020).

## 5 Conclusion

We acquired, for the first time, in a systematic large-scale area-wide survey presence/absence data of (a total of 5,160) red wood ant nests (RWA; *Formica polyctena*) in two study areas in the Oberpfalz, NE Bavaria, Germany. We investigated both, whether variable factors such as physical nest parameters, tree species and age, nest building material and quasi-invariant factors, e.g. geochemical composition of bedrock, geomorphology, exposure and tectonics have an influence on the spatial distribution and nest densities of RWA nests. A combination of mature (≥80–140 years) pine-dominated forests as main variable factor and the geochemical property of the Variscan granites with its high natural Rn potential and the moderate heat production of the Variscan granites as main quasi-invariant factor could explain the high nest numbers at both sites. The spatially clustered distribution patterns of the observed nests suggest a strong interaction between nests and their quasi-invariant environment, especially the directionality of the present-day stress field and the direction of the tectonically formed “Erbendorfer Line”. In general, such a combination of variable and quasi-invariant factors can be addressed as particularly favorable ant habitats. Future investigations will show, whether high nest numbers can be observed in comparable geological and geochemical environments, e.g. in other areas with Variscan granites with similar high Rn potential and the moderate heat production. We suggest that favorable geochemical and tectonic factors, which have been neglected so far, need to be considered in future assessments of RWA nests occurrences.

## 6 Acknowledgements

The field work and first analyses were conducted during the time, the first and corresponding author (Gabriele M. Berberich) was research associate of the Technical University of Dortmund. G.M.B’s field work was funded by Bayerische Staatsforsten, Regensburg (grant number 19/Wa/N6). We greatly acknowledge the Landesamt für Digitalisierung, Breitband und Vermessung (LDBV), München, that provided the 1 m DTM for terrain analyses and the Bayerische Staatsforsten (BSF) AöR, Regensburg, that provided data from the 10-year forest inventory and management plan of 2009. Either the Landesamt für Digitalisierung, Breitband und Vermessung (LDBV), München nor the Bayerische Staatsforsten had a role in the design of the study; in the collection, analyses, or interpretation of data; in the writing of the manuscript, and in the decision to publish the results. We greatly acknowledge A. M. Ellison, Harvard Forest, Harvard University for reviewing the draft.

## Author contributions

G.M.B. conceived the idea, designed the study, performed the field work, carried out the statistical analysis and wrote the manuscript. M.B.B. performed the field work, analyzed the data and contributed to the manuscript. M.G. contributed to the manuscript. All authors edited the manuscript and approved the final version.

